## Supplementary material for "Characterization of humoral and SARS-CoV-2 specific T cell responses in people living with HIV": Suppl. Table S2

**Fluorochrome-Conjugated Antibodies:**

| Antibodies | Supplier | Identifier | Clone |
|---|---|---|---|
| APC anti-human IgM Antibody | BioLegend | Cat # 314510 | Clone # MHM-88 |
| APC/Cyanine7 anti-human CD19 Antibody | BioLegend | Cat # 363010 | Clone # SJ25C1 |
| APC/Cy7 anti-human CD197 (CCR7) | BioLegend | Cat # 353212 | Clone # G043H7 |
| Brilliant Violet 605™ anti-human CD3 Antibody | BioLegend | Cat # 317322 | Clone # OKT3 |
| Brilliant Violet 650™ anti-human CD127 (IL-7Rα) Antibody | BioLegend | Cat # 351325 | Clone # A019D5 |
| Brilliant Violet 650™ anti-human CD3 Antibody | BioLegend | Cat # 317324 | Clone # OKT3 |
| Brilliant Violet 711™ anti-human CD27 Antibody | BioLegend | Cat # 302833 | Clone # O323 |
| Brilliant Violet 785™ anti-human CD38 Antibody | BioLegend | Cat # 303530 | Clone # HIT2 |
| Alexa Fluor® 700 anti-human CD45RA Antibody | BioLegend | Cat # 304120 | Clone # HI100 |
| PE/Cyanine7 anti-human CD45RA Antibody | BioLegend | Cat # 304126 | Clone # HI100 |
| Brilliant Violet 421™ anti-human CD279 (PD-1) Antibody | BioLegend | Cat # 329920 | Clone # EH12.2H7 |
| PE/Dazzle™ 594 anti-human CD4 Antibody | BioLegend | Cat # 300548 | Clone # RPA-T4 |
| PE anti-human IgD Antibody | BioLegend | Cat # 348204 | Clone # IA6-2 |
| APC anti-human IFN-γ Antibody | BioLegend | Cat # 506510 | Clone # B27 |
| Brilliant Violet 785™ anti-human CD8a Antibody | BioLegend | Cat # 301046 | Clone # RPA-T8 |
| Brilliant Violet 711™ anti-human CD8a Antibody | BioLegend | Cat # 301044 | Clone # RPA-T8 |
| Brilliant Violet 510™ anti-human CD4 Antibody | BioLegend | Cat # 300546 | Clone # RPA-T4 |
| Brilliant Violet 510™ anti-human CD14 Antibody | BioLegend | Cat # 301842 | Clone # M5E2 |
| Brilliant Violet 510™ anti-human CD19 Antibody | BioLegend | Cat # 302242 | Clone # HIB19 |
| Brilliant Violet 711™ anti-human CD279 (PD-1) Antibody | BioLegend | Cat # 329928 | Clone # EH12.2H7 |
| Brilliant Violet 711™ anti-human PD-1 Antibody | BioLegend | Cat # 300232 | Clone # RPA-2.10 |
| PE/Cyanine7 anti-human CD154 Antibody | BioLegend | Cat # 310832 | Clone # 24-31 |
| APC-R700 Mouse Anti-Human CD196 (CCR6) | BD Biosciences | Cat # 565173 | Clone # 11A9 |

| | | | |
|---|---|---|---|
| BB515 Rat Anti-Human CXCR5 (CD185) | BD Biosciences | Cat # 564624 | Clone # RF8B2 |
| BV605 Mouse Anti-Human CD56 | BD Biosciences | Cat # 562780 | Clone # NCAM16.2 |
| PC-Cy7 Mouse Anti-Human CD25 | BD Biosciences | Cat # 335824 | Clone # 2A3 |
| BB700 Mouse Anti-Human CD16 | BD Biosciences | Cat # 746199 | Clone # 3G8 |
| BB700 Mouse Anti-Human CD4 | BD Biosciences | Cat # 566393 | Clone # SK3 |
| PE-Cy™5 Mouse Anti-Human CD183 | BD Biosciences | Cat # 551128 | Clone # 1C6/CXCR3 |
| PE-Cy™5 Mouse Anti-Human HLA-DR | BD Biosciences | Cat # 562007 | Clone # G46-6 |
| FITC Mouse Anti-Human TNF-α | BD Biosciences | Cat # 554512 | Clone # MAb11 |
| GolgiStop (with Monensin) | BD Biosciences | Cat # 554724 | |
| PerCP-eFluor 710 Anti-Human IL-2 | eBioscience | Cat # 46-7029-42 | Clone # MQ1-17H12 |
| PerCP-eFluor 710 Anti-Human CD3 | eBioscience | Cat # 46-0037-42 | Clone # OKT3 |
| PE Anti-Human TIGIT | eBioscience | Cat # 12-9500-42 | Clone # MBSA43 |

**Key Chemicals, Peptides, and Commercial Assays**

| Reagents | Supplier | Identifier |
|---|---|---|
| PepTivator SARS-CoV-2 Prot_N | Miltenyi Biotec | Cat # 130-126-698 |
| PepTivator SARS-CoV-2 Prot_M | Miltenyi Biotec | Cat # 130-126-702 |
| PepTivator CMV pp65, human | Miltenyi Biotec | Cat # 130-093-438 |
| ProMixTM HIV Peptide Pool | Proimmune | Cat # PX-HIV |
| Human IFN-γ ELISpot Kit | Mabtech | Cat # P3420-2A |
| Human Anti-Cytomegalovirus IgG ELISA Kit (CMV) | Abcam | Cat # ab108724 |
| Brefeldin A | eBioscience | Cat # 00-4506-51 |
| Foxp3/TF Staining Buffer Set | Invitrogen | Cat # 00-5523-00 |
| BD Cytofix/Cytoperm™ Fixation/Permeabilization Solution Kit | BD Biosciences | Cat # 554714 |
